# Delineating the functional role of the *PPE50 (Rv3135) - PPE51 (Rv3136)* gene cluster in the pathophysiology of *Mycobacterium tuberculosis*

**DOI:** 10.1101/2023.03.30.534876

**Authors:** Ravi Prasad Mukku, Kokavalla Poornima, Sangya Yadav, Tirumalai R. Raghunand

## Abstract

The extraordinary success of *Mycobacterium tuberculosis* (*M. tb*) has been attributed to its ability to modulate host immune responses. The genome of *M. tb* encodes multiple immunomodulatory factors, including several proteins of the multigenic PE_PPE family, which comprise about 10% of its coding potential. The presence of these proteins in pathogenic mycobacteria strongly suggests that they play a role in disease pathogenesis. To understand its role in *M. tb* physiology we have begun to characterise the *PPE50 (Rv3135)*-*PPE51 (Rv3136)* gene cluster, one of nine *PPE-PPE* clusters in the *M. tb* genome. We demonstrate that this cluster encodes a co-transcriptional unit and that PPE50 and PPE51 interact both *in vitro* and *in vivo*, the first demonstration of PPE-PPE interaction. THP-1 macrophages infected with recombinant *M. smegmatis* strains expressing *PPE50* and *PPE51* showed less intracellular viability than the control strain containing the vector alone, the decline in viable counts correlating with an increase in transcript levels of inducible nitric oxide synthase (*iNOS2*). Macrophages infected with the recombinant strains exhibited an upregulation in levels of the anti-inflammatory cytokine *IL-10*, indicating an immunomodulatory role for these proteins. Using pull-down assays, we discovered TLR1 to be the cognate receptor for PPE50, with signalling through the receptor being indicated by an increase in IRAK1 phosphorylation. All the phenotypes observed on infection of THP-1 macrophages including the decrease in CFUs, the increase in *iNOS2* and *IL-10* levels, as well as signalling through the receptor, were reversed on treatment of macrophages with an anti-TLR1 antibody prior to infection, validating the functional outcome of PPE50-TLR1 interaction. Our data points to a TLR1 dependent role for the *PPE50-PPE51* cluster in promoting bacillary persistence, *via* CFU reduction and a concomitant upregulation of the anti-inflammatory response - a two-pronged strategy to circumvent host immune surveillance.

## INTRODUCTION

*M. tb*, the causative agent of human tuberculosis, has achieved unprecedented success due to its ability to modify host immunological responses, thereby promoting its long-term survival (1). Understanding the pathogenic processes of *M. tb* depends on identifying and characterizing bacillary elements involved in immune evasion (2), and their interaction with host defense components during infection. In this context, the multigenic PE/PPE proteins, which account for 10% of the coding potential of the *M. tb* genome, have emerged as key players because several family members have been linked to evasion of host immune processes. These proteins are named after the conserved Proline-Glutamate and Proline-Proline-Glutamate residues at their N-termini (3, 4), and the *M. tb* genome is reported to encode 169 PE/PPE proteins. A wide variety of *PE/PPE* genes have been discovered within pathogenic Mycobacteria. Most of them are arranged sporadically in the genome, and a few are arranged in clusters. *M. tb* has total of 40 such *PE*/*PPE* gene clusters, with loci including *PE-PPE* [15], *PE-PE* [5] and *PPE-PPE* [9] genes arrayed in tandem; among them are uncommon clusters like a *PPE* gene followed by another *PPE*, and a *PE* gene sandwiched between two *PPE* genes (3). Putative operonic clusters containing only *PE*/*PPE* genes are found for 22 of the 69 PPE proteins (3, 5). The 14 gene pairs with a 5’ *PE-PPE* 3’ orientation are separated by 100 bases or less, and are likely to be co-transcribed and be functionally linked (6). Evidence from comparative genomics suggests that the expansion of the *PE*/*PPE* family is linked to the five *Esx* clusters spread out over the *M. tb* genome, which encode components of the Type VII secretion system. In contrast to the extensive studies on the PE_PGRS sub-group in the literature, no studies have been conducted on the role of *PPE-PPE* clusters in *M. tb* pathophysiology. We have selected the *Rv3135* (*PPE50*) - *Rv3136* (*PPE51*) cluster for its functional characterisation and to understand the involvement of the two genes in *M. tb* physiology, owing to their peculiar cluster arrangement. In a microarray study, these genes were observed to be upregulated on exposure to palmitic acid. Repression in expression of both genes was seen on exposure of *M. tb* to DETA/NO, H_2_O_2_, KCN, and under stationary phase growth conditions (7). More recent reports have suggested a role for PPE51 in drug tolerance (8), nutrient transport (9), and in the inhibition of autophagy (10). No functional studies have yet been described for the individual genes, or for the gene cluster as a whole.

## MATERIALS AND METHODS

### Bacterial strains, media and growth conditions

*M. smegmatis* mc^2^6 and *M. tb* H37Rv were cultivated in Middlebrook, 7H9 broth and 7H10 agar (Difco) containing albumin dextrose complex (5 g BSA, 2 g glucose, and 0.85 g NaCl/L), 0.5% (v/v) glycerol, and 0.5% (v/v) Tween 80. Luria Bertani medium was used to cultivate *E. coli* strains - both *E. coli* and mycobacteria were cultured by shaking at 37° C. When required, 200 µg/ml of ampicillin and 50 µg/ml of kanamycin (for *E. coli*) and 15 µg/ml of kanamycin (for mycobacteria) were added to the culture medium.

### DNA techniques

All restriction enzymes, T4 DNA ligase, and Taq polymerase were purchased from New England Biolabs (NEB) and Invitrogen, respectively. DNA manipulation protocols including plasmid DNA preparation, restriction endonuclease digestion, agarose gel electrophoresis, ligation of DNA fragments and *E. coli* transformation were performed according to (11). The PCR amplifications were performed according to the manufacturer’s instructions. During each cycle, 95° C was held for 30 s, 60° C for 30 s, and 72° C for 1 min, followed by a 72° C extension cycle for 10 min. The Qiagen gel extraction kit was used to purify DNA fragments for cloning reactions according to the manufacturer’s instructions. *M. smegmatis* was transformed by electroporation.

### In silico analyses

Multiple Sequence Alignments were performed using the ClustalW2 algorithm(12). All *M. tb* sequences were obtained from the Mycobrowser database (https://mycobrowser.epfl.ch). The TM-pred (13) and Kyte& Doolittle (14) algorithms were used to identify trans-membrane domains and regions of hydrophobicity respectively.

### Sub-cellular localisation and Proteinase K Sensitivity Assay

In order to determine the subcellular location of these proteins, the *M. tb PPE50*, *PPE51* ORFs were cloned between *BamHI* and *EcoRI* sites of pJEX55 (15) and the recombinant plasmids were transformed into *M. smegmatis*. The recombinant *M. smegmatis* strains expressing c-myc tagged PPE50, PPE51 were harvested at the logarithmic phase of growth, washed and resuspended in PBS. The samples were divided into two identical aliquots and incubated at 37°C for 30 min with or without Proteinase K (Sigma). With the addition of 2 mM EGTA, the reaction was stopped, and the sub-cellular fractions of these samples were isolated as described in (16). By denaturing SDS PAGE, the individual fractions were separated, and fusion proteins were detected by western blotting with anti-c-myc antibodies (sc40, Santa Cruz).

### Expression of PPE50, PPE51 in *M. smegmatis*

*M. tb PPE50, PPE51* ORFs were amplified using gene-specific primers (Table S1) and cloned between the *BamHI* and *EcoRI* sites of pMV261(17) These constructs were transformed into *M. smegmatis* for overexpression.

### Expression and purification of PPE50 and PPE51

The full-length *M. tb PPE50* ORF was cloned as C-terminal 6xHis-tagged fusion in pET22b, and *PPE51* was cloned as N-terminal Glutathione-S-Transferase (GST) tagged fusion in pGEX6p1. Following sequence confirmation, the constructs were transformed into *E. coli* BL21(DE3). For induction, cultures were allowed to grow to an OD of 0.5, induced with 0.5 mM IPTGm following which PPE50 was purified using Ni-NTA affinity chromatography, and PPE51 was purified using glutathione agarose beads. Purified proteins were used for pull-down experiments.

### *In vitro* growth kinetics

To functionally characterise *M. tb PPE50, PPE51* their ORFs were amplified from *M. tb* H37Rv genomic DNA using gene specific primers (Table S1), cloned between the *BamHI* and *EcoRI* sites of pMV261 (18) and transformed into *M.smegmatis*. To examine their growth patterns, recombinant *M. smegmatis* strains were grown until late exponential phase, diluted to an A_600_ (OD) of 0.2 and cultured in Middlebrook 7H9 broth containing 15 µg/ml kanamycin. Growth curves were generated by CFU measurements and plotted against time. At each designated time point, cultures were harvested for RNA extraction and gene expression analyses performed. All CFU enumeration assays were performed in the presence of 15 μg/ml kanamycin.

### Macrophage infection

THP-1 monocytes were cultured at 37° C in 5% CO_2_ in RPMI 1640 medium supplemented with 10% (v/v) Fetal bovine serum and antibiotics (60 µg/ml penicillin G sodium, 50 µg/ml streptomycin sulphate, and 30 µg/ml gentamycin sulphate). For the infection experiments, monocytes were seeded at a density of 2×10^6^/well for 24 h, differentiated with 5 ng/ml PMA, and then infected 72 h later. Cultures of exponentially growing bacteria were pelleted, washed, resuspended in RPMI medium (without antibiotics) to an OD of 1.0. Single cell suspensions of recombinant *M. smegmatis* strains were prepared by passing culture suspensions through needles of 26 1/2 gauge 5-6 times. At each step, CFU counts were performed to determine bacterial viability. Based on pilot infections with multiple MOIs we performed with the cell lines used, equal numbers of each strain were used for infections (Input counts) at a MOI of 1:100. Post-infection CFU counts were determined by lysis of infected cells after 2 h of incubation with bacteria (T0 counts). In order to kill extracellular bacteria, complete RPMI containing gentamycin was added. Infected cells were lysed with 0.1% Triton X-100 and plated on Middlebrook 7H10 agar at the designated time points to determine CFU counts. For each experiment, macrophages infected with *M. smegmatis* expressing the empty vector were used as controls.

### Real-time RT-PCR analysis

The expression profiles of *PPE50*, *PPE51* in *M. tb* and recombinant strains of *M. smegmatis* expressing pMV261-*PPE50*, pMV261-*PPE51* as a function of growth, were determined using real-time transcript measurements. For this, cells were harvested at the OD 0.8, 1.5, 2, 3, 3.5 and at 4, 8, 12, 24, 48 and 72 h time points respectively, and total RNA isolated from each culture using TRIzol reagent (Invitrogen) as per the manufacturer’s protocol. Following treatment with RNAse free DNAse I, cDNA synthesis was performed using the iScript cDNA synthesis kit (Bio-Rad) and subsequently used as a template for SYBR green based PCR amplification using *PPE50*, *PPE51* specific primers (Table S1) to generate 200 bp amplicons. Gene specific transcript levels in *M. smegmatis* were normalised to the *sigA* transcript in each sample and in *M. tb* to the 16s rRNA transcript. In *M. smegmatis* the relative fold change in transcript levels at each time point was calculated with respect to the levels at 4 h. In the *M. tb* expression profile, the relative fold change in transcript at each OD point was calculated with respect to the levels at OD 0.8. To quantify host cytokines and *iNOS2* transcript levels, total RNA was isolated from infected macrophages using TRIzol reagent. Following treatment with RNAse free DNAse I, cDNA synthesis was performed using the iScript cDNA synthesis kit (Bio-Rad) and subsequently used as a template for SYBR green based PCR amplification using primers specific to *iNOS*, *IL-10*, *IL-12* (Table S1) designed to generate 200 bp amplicons. The levels of each mRNA were normalised to the transcript levels of GAPDH and β-actin. Relative fold change was calculated with reference to macrophages infected with *M. smegmatis* expressing the empty vector.

### Receptor interaction assay

To extract the membrane fraction for receptor identification, PMA-differentiated THP-1 cells were resuspended in cell lysis buffer (50 mM HEPES buffer pH 7.4, 150 mM NaCl, 5 mM EDTA, PIC), sonicated at 20 percent amplitude with 10 s on and off for 5 minutes, centrifuged at 13000 rpm for 1 h, and the pellet was resuspended in lysis buffer containing 1% CHAPS. For co-immunoprecipitation assays, 50 µg of PPE50-6xHis was combined with 500 µg of cell membrane fractions in lysis buffer to a final volume of 500 µl. The reactions were incubated at 37° C for 6 h, after which anti-His antibodies (CST) were used to conduct pull-downs. Before adding antibodies, 60 µl of each sample was aliquoted for use as loading controls. Each sample was then mixed with Protein A agarose beads and incubated overnight at 4° C. Following this, after three washes with PBS, the beads were boiled in 50 µl of sample buffer, resolved on SDS-PAGE. Following transfer onto PVDF membranes, the co-immunoprecipitated proteins were identified using antibodies to TLR1/2/ 4/ 6. Equal loading of the cell membrane fraction was evaluated using western blotting using antibodies to CD81. In reverse pulldown assays we incubated PPE50 with the membrane fraction along with a TLR1 Ab (Abcam) at 4^0^ C overnight, bound the mix with Protein A beads, washed and resolved the bead bound proteins on SDS-PAGE and transferred them onto PVDF membranes. The blot was then probed with an anti-His antibody (CST).

### TLR1 Receptor inhibition

THP-1 monocytes were grown in RPMI 1640 media supplemented with 10% (v/v) Fetal bovine serum and antibiotics (60 µg/ml penicillin G sodium, 50 µg/ml streptomycin sulphate, and 30 µg/ml gentamycin sulphate) at 37°C and 5% CO_2_. These monocytes were seeded at a density of 2×10^6^ per well for 24 h, differentiated with 5 ng/ml PMA for 72 h, and then infected. Prior to infection, we treated these differentiated cells for 6 with 0.5µg/ml of a polyclonal anti-TLR1 antibody (Abcam).

### Western blotting

The levels of total and phosphorylated IRAK-1 in infected THP-1 macrophages were determined by western blot analysis. Each sample was lysed in RIPA lysis buffer, following which, equal protein amounts from each cell lysate subjected to SDS-PAGE and transferred onto PVDF membranes. Each blot was sequentially probed with antibodies specific to phosphorylated or total IRAK1 (Cell Signalling Technology & Abcam) and β-actin. Proteins were detected using Enhanced Chemiluminescence (Santa Cruz). For the inhibition studies, THP-1 cells were treated with 0.5 µg/ml anti-TLR1 Ab (Abcam) 6 h prior to infection. Recombinant *M. smegmatis* containing pMV261 was used as an empty vector/ mock control. For quantitation, the intensity of all bands was densitometrically measured using ImageJ and normalised to levels of β-actin. The normalised levels of phosphorylated IRAK1 in control samples (pMV261) were assigned a value of 1, and the fold change in the levels of these proteins in the test samples (PPE50, PPE51) were derived with respect to the control.

### Co-transcriptional analysis

Total RNA extracted from exponentially growing *M. tb* H37Rv was used to assess the transcriptional state of the *PPE50-PPE51* cluster. RNA treated with DNAse I was used as a template for cDNA synthesis with the primers *PPE51* R1 (Table S1). The cDNA was utilised for PCR amplification with gene-specific primer combinations. The analysis includes appropriate negative controls (-RT).

### Co-immunoprecipitation

To perform *in vivo* pull down assays, 150 ml of independent *M. smegmatis* cultures co-transformed with pJEX55+pSCW54, pJEX55-*PPE51*+pSCW54, pJEX55+pSCW54-*PPE50*, pJEX55-*PPE51*+pSCW54-*PPE50* were grown in Middlebrook 7H9 broth containing kanamycin and hygromycin to log phase and induced with 0.2% acetamide for 6 h. Induced cells were pelleted, resuspended in 1.2 ml PBS containing protease inhibitors and lysed by sonication. Half of this lysate was used to prepare a cell envelope fraction by ultracentrifugation (16) and the pellets were resuspended in 150 µl of PBS containing protease inhibitors and 0.5% CHAPS. 400 µl of cell lysates were mixed with 50 µl of the solubilised cell envelope fraction and the protein amounts in the mix were quantitated using a Bradford assay. For the co-IP assay 500 µg of protein in a total volume of 500 µl PBS was incubated with an anti c-myc antibody; prior to antibody addition, 30 µl of the reaction mix was removed and saved as a loading control. Protein A agarose beads were then added to each sample and incubated at 4°C for 6 h. Following three washes with PBS, the beads were boiled in 50 µl of sample buffer and subjected to SDS-PAGE. Separated proteins were transferred to PVDF and probed with an anti-His antibody.

### Mycobacterial Protein Fragment Complementation (MPFC) assay

To assess the interaction of PPE50-PPE51 at *in vivo PPE50*, *PPE51* were cloned into the *E. coli* - *Mycobacterium* shuttle vectors pUAB400 and pUAB300 respectively. Both constructs were co-transformed in *M. smegmatis* mc^2^155 and the interaction assay was performed as described (19). Appropriate positive (pUAB100+pUAB200) and negative (pUAB100+pUAB400, pUAB400+ pUAB300) controls were included in the assay.

## RESULTS

### *PPE50* (*Rv3135*) and *PPE51* (*Rv3136*) are co-operonic and interact with each other

Among the 40 PE/PPE clusters in the *M. tb* genome, *PPE50-PPE51* is one of 9 such PPE-PPE clusters. The two proteins share 62% identity (71/114) and 77% homology (88/114) in their PPE domains with PPE51 (380 aa) possessing a C-terminal extension that PPE50 (132 aa) lacks (Fig. 1). Since *PPE50-PPE51* was among the nine *PPE-PPE* gene pairs expected to be co-operonic based on their co-repression in drug tolerance research, it was hypothesised that they would be cooperative (8). To confirm this hypothesis, RT-PCR analyses were performed with RNA isolated from exponentially growing *M. tb* H37Rv using primer pairs (Table S1) designed to amplify ORF and junction-specific areas of the gene pair (Fig. 2A) (*PPE50*-200 bp, *PPE51*-200 bp, and *PPE50-PPE51* junction-200 bp). We observed amplification products in the test lanes but not in the-RT control lanes, leading us to infer that this gene pair was transcribed as a mono-cistronic message (Fig. 2B). Since co-operonic protein pairs are known to often physically interact (20–22) the potential interaction of the PPE50-PPE51 protein pair was investigated by performing co-immunoprecipitation of C-terminally His-tagged PPE50 with N-terminally GST-tagged PPE50 (Fig. S1) along with the appropriate controls. Co-precipitation of PPE51 and PPE50 suggested these two proteins directly interact *in vitro* (Fig. 2C). To verify this finding, *in vivo* pull-down experiments were performed utilising cell wall enriched cell lysates of *M. smegmatis* expressing C-terminal c-myc fused PPE51 and C-terminal 6xHis-tagged PPE50. Co-precipitation of PPE50 and PPE51 suggested that the two proteins interact *in vivo* as well (Fig. 2D). We ensured that in both the *in vitro* and *in vivo* pull-down analyses, equal amounts of each fraction were loaded (Fig. S2). In addition, the mycobacterial protein fragment complementation (M-PFC) assay was used to examine the interaction between these proteins (19). Co-transformants of *M. smegmatis* mc^2^155 expressing two protein pairs fused to the murine dihydrofolate reductase domains F [1,2] and F [3] were resistant to 40 g/ml trimethoprim (Figure 2 E). Negative control samples exhibited no growth, validating the *in vivo* interaction between PPE50 and PPE51.

**Figure 1:**
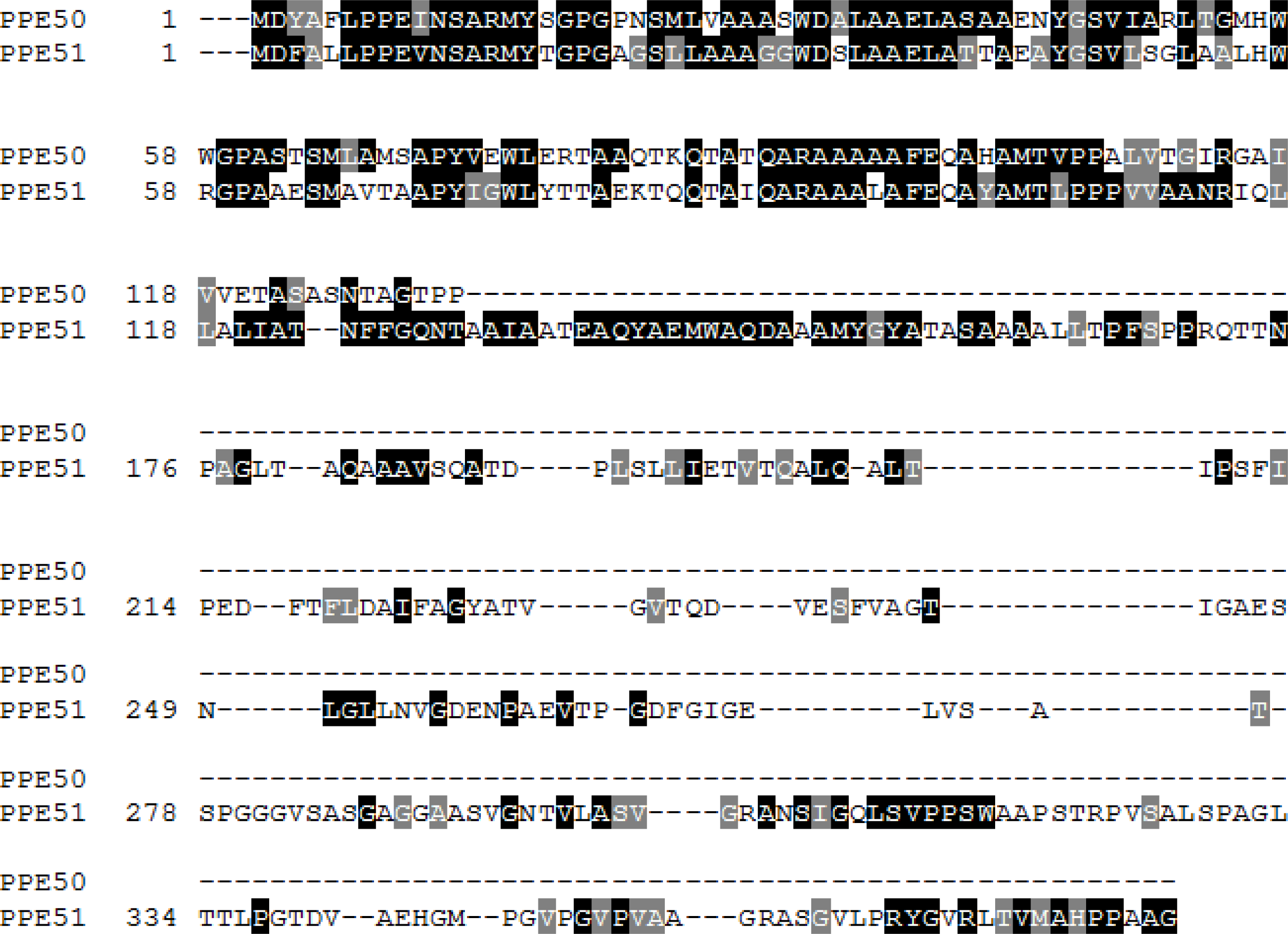
Sequence homology of *M. tb* PPE50/PPE51. Boxshade representation of PPE50, PPE51 protein sequence alignment.

**Figure 2:**
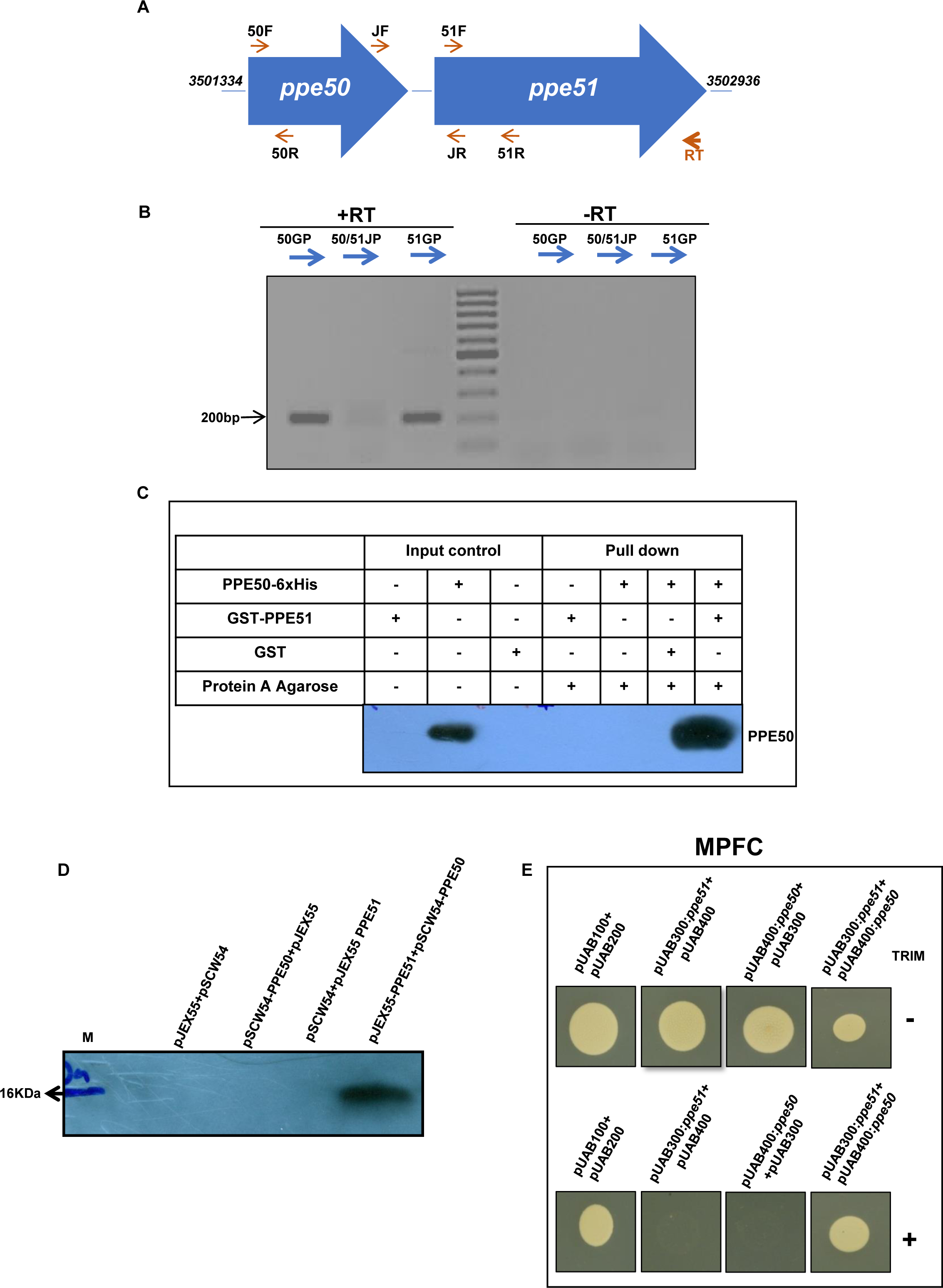
*PPE50* and *PPE51* are co-transcribed, and their gene products interact. (A) Schematic depiction of the genomic structure of the *PPE50-PPE51* region in the *M. tb* genome showing the sites of the primers used in the investigation. (B) RT-PCR amplification products of the *PPE50-PPE51* gene pair: Lane 1-(50F+50R), Lane 2-(50JF+51JR), Lane 3 - (51F+51R), Lanes 5, 6, and 7 correspond to-RT controls for these respective primer pairs, Lane 4-100 bp DNA ladder. (C) Western blot of *in vitro* co-immunoprecipitation reactions of PPE50-6xHis and GST-PPE51 pulled with an anti-GST antibody and probed with anti-His antibody. (D) Western blot of PPE50-PPE51 co-IP from *M. smegmatis* expressing PPE51-myc and PPE50-6xHis. The complex was pulled down using an anti-c-myc antibody, and the blot was probed with an anti-His antibody. (E) M-PFC analysis of the interaction of PPE50 and PPE51. The image depicts the growth pattern of *M. smegmatis* co-transformants on Middlebrook 7H11 agar plates with (+) or without (-) 40µg/ml trimethoprim (TRIM). The findings are based on at least two biological replicates.

### *M. smegmatis* strains expressing *PPE50* and *PPE51* show reduced survival in macrophages

To investigate the possible involvement of PPE50 and PPE51 in mycobacterial pathophysiology, we used the saprophyte *M. smegmatis,* which has been extensively used as a surrogate model to study the roles of this family of proteins, including by our own research group (20, 21, 23–27). Since it lacks the homologues of most PE-PPE proteins, including *PPE50* and PPE51, we expressed these proteins in *M. smegmatis* and examined their localisation. To do so, recombinant *M. smegmatis* strains expressing c-myc tagged PPE50 and PPE51 were treated with Proteinase K, and sub-cellular fractionation was carried out. Prior to western blotting with a c-myc antibody, we ensured equal loading for each fraction (Fig. S3A, B). The blots revealed that both proteins were localized to the cell wall and, that their signals were greatly diminished in the cell wall-associated fractions treated with Proteinase K (Fig. S34, B). This indicated that PPE50 and PPE51 are exposed to the cell surface, making this a suitable system to evaluate their functions as modulators of pathogenicity *via* contact with the host. This finding was consistent with the predicted hydrophobicity and transmembrane domain prediction of these proteins generated *in silico* (Fig. S5A, B). *M. smegmatis* strains expressing untagged PPE50 and PPE51 under the control of the constitutive *hsp60* promoter were then used to determine whether these proteins mediate host-pathogen interactions (17, 18). CFU enumeration and growth profiles (Fig. S5A, B) of recombinant *M. smegmatis* strains expressing *PPE50*, *PPE51*, *PPE50-PPE51*, and the empty vector revealed that over-expression of these genes did not result in growth compromise *in vitro*. Both genes were expressed at comparable levels in these strains as quantified by real-time RT-PCR using gene-specific primers (Table S1), suggesting that their expression did not vary as a function of growth (Fig. S6C, D, E). Using the THP-1 macrophage infection model, we assessed the intracellular survival of the recombinant *M. smegmatis* strains, and the ability of the proteins to modulate host functions. Curiously, bacillary counts of strains expressing *PPE50*, *PPE51*, and *PPE50-PPE51* were lower than those of the empty vector control at 24, 48, and 72 hours post-infection in THP-1 macrophages (Fig. 3A), Though unexpected, this observation correlated with the elevated levels of inducible nitric oxide synthase-2 (*iNOS2*) expression in these cells (Fig. 3B). This is significant since iNOS2 is an important factor in the control of intracellular bacillary survival (28–30).

**Figure 3:**
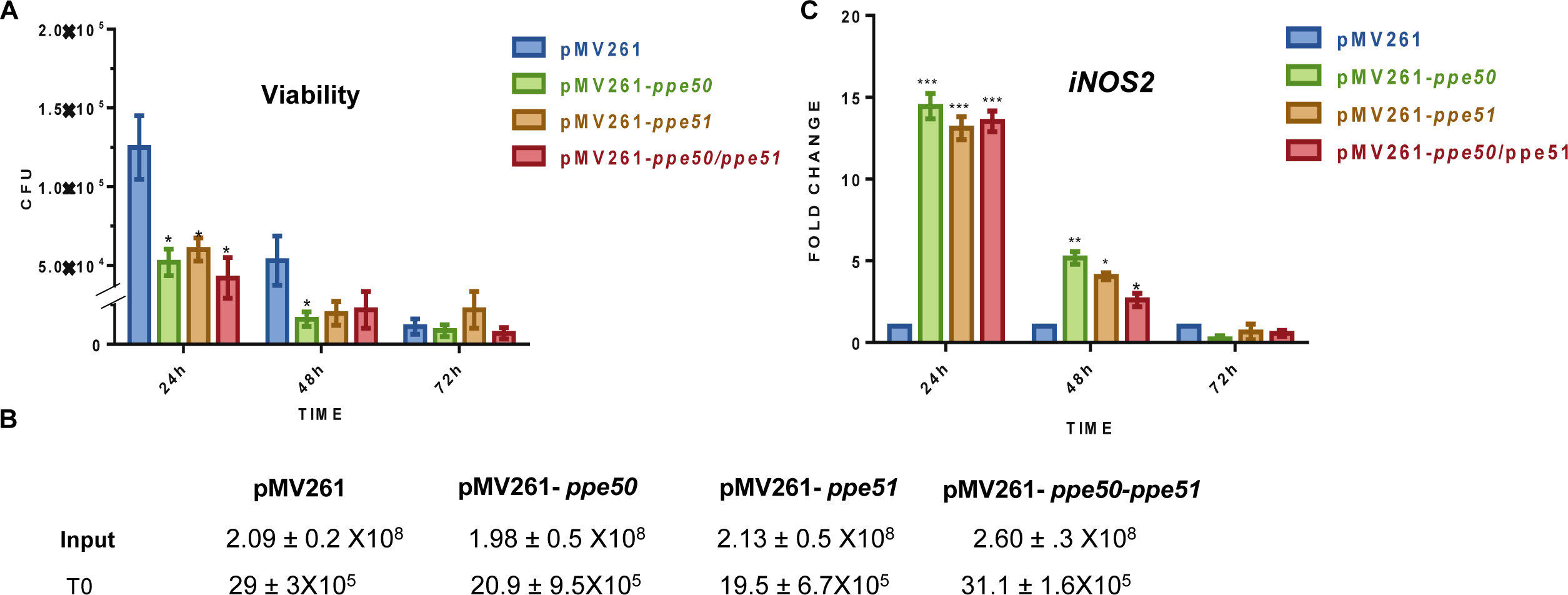
Intra macrophage growth of recombinant *M. smegmatis* expressing *PPE50* and *PPE51*. CFU counts of *M. smegmatis* expressing pMV261, *PPE50*, *PPE51* and *PPE50-PPE51*, 24 h, 48 h and 72 h post infection in THP-1 macrophages (A) Input and T_0_ (post-infection) CFU counts of infecting bacilli (± SEM) (B). *iNOS2* transcript levels in THP-1 macrophages infected with *M. smegmatis* expressing pMV261, *PPE50*, *PPE51* and *PPE50-PPE51*, 24 h, 48 h and72 h post infection (C). The data is representative of two biological replicates. *p 0.05, **p 0.005, ***p 0.001 Error bars represent mean ± SEM of three ≤ ≤ biological replicates.

### PPE50 and PPE51 are immunomodulators that promote IL-10 production through TLR1 signalling

Our observation that PPE50 and PPE51 are engaged in reducing intra-macrophage bacterial survival, led us to explore if they also affect immunological responses, as has been observed for other members of this protein family (5, 20, 21, 25, 31). To test their potential immune modulatory function, we infected THP-1 macrophage with recombinant *M. smegmatis* strains expressing *PPE50*, *PPE51* and *PPE50-PPE51*, and examined the levels of pro-and anti-inflammatory cytokines in infected cells. As shown in Fig. 4A, we observed an upregulation of the anti-inflammatory cytokine IL-10 in macrophages infected with recombinant *M. smegmatis* expressing *PPE50* at 24 h,48 h and 72 h post-infection, and the strain expressing *PPE50-PPE51* at the 24 h time point, in comparison to controls. Working on the hypothesis that this induction must happen *via* ligand-receptor interaction, we used purified PPE50 and PPE51 (Fig. S1) to identify their cognate receptors. In pull-down and reverse pull-down experiments with macrophage membrane fractions, we identified TLR1 to be the binding partner for PPE50 (Fig. 4B, C); in the same experiments, no TLR interaction could be identified for PPE51 (data not shown). Equal loading was ensured for all fractions in these experiments, by quantifying levels of CD81 (Fig. S7). To test the functionality of PPE50:TLR1 interaction, we monitored IRAK1 phosphorylation as a marker for induction of TLR1 signalling (32), in macrophages infected with the above-listed recombinant *M. smegmatis* strains. We observed an increase in phosphorylation of IRAK-1 in THP-1 macrophages infected with the recombinant PPE50-expressing *M. smegmatis* strain, implying a functional consequence for PPE50:TLR1 interaction; no induction of IRAK-1 phosphorylation was seen in the other two infection samples (Fig. 4D, E). To test if the induction of *iNOS2* and *IL-10* we observed in earlier experiments was mediated by PPE50:TLR1 interaction, we performed TLR1 blocking experiments by incubating macrophages with an anti-TLR1 antibody, prior to infection with the *PPE50* expressing recombinant *M. smegmatis* strain. As depicted in figure 5, all the phenotypes associated with infection of macrophages with recombinant *M. smegmatis* expressing *PPE50*, were reversed on TLR1 blocking compared to the controls. These included the reduction in viability (Figure 5A, B), the increase in *iNOS2* transcripts levels (Figure 5C, D), as well as the elevation in the *IL-10* transcript (Figure 5E, F). Moreover, the increase in IRAK1 phosphorylation was lost on antibody blocking as well (Figure 9G, H), clearly indicating that PPE50 activates the IRAK1 pathway through TLR1 to upregulate the expression of *iNOS2* and *IL-10*.

**Figure 4:**
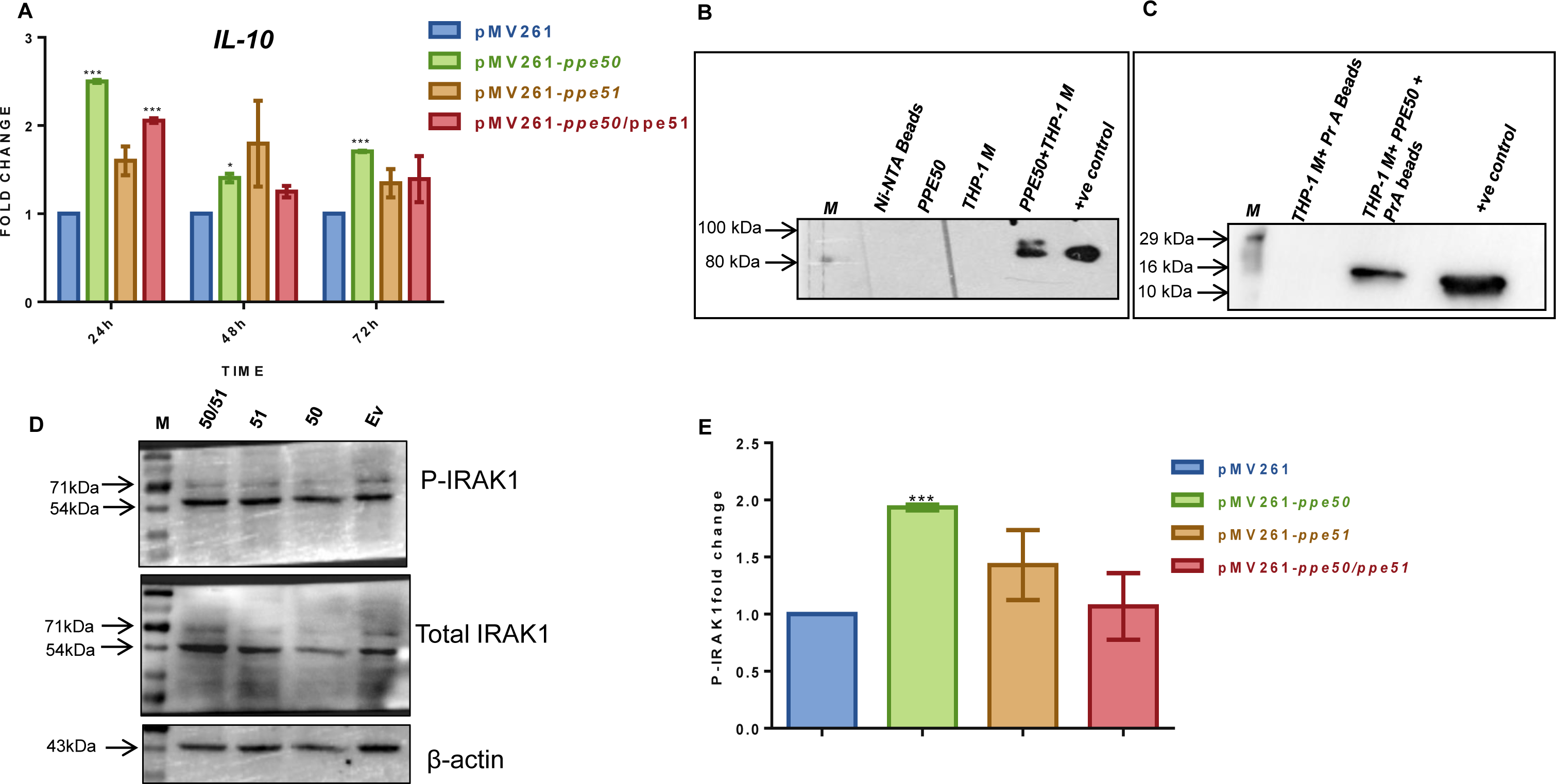
Immunomodulatory effects of the PPE50-PPE51 cluster. (A) *IL-10* transcript levels following THP-1 infection with recombinant *M. smegmatis* strains expressing *PPE50*, *PPE51*, *PPE50PPE51*,and the empty vector pMV261. (B) Receptor recognition by PPE50 Western blot depicting co-IP reactions of PPE50-His and solubilised cell-membrane fractions of THP-1 macrophages-an anti-His antibody was used pull-down the complex and the blot was probed with an anti-TLR1 Ab. (C) Reverse pull-down for receptor recognition – an anti-TLR1 antibody was used to pull down the complex and the blot was probed with an anti-His antibody. (D) Western blots showing levels of phospho-IRAK1, Total IRAK1 and β-actin levels in lysates from Macrophages infected with recombinant *M. smegmatis* strains expressing *PPE50* (50), *PPE51* (51), *PPE50-PPE51* (50/51) and the empty vector pMV261 (EV). The densitometric quantitation of this data is represented in (E). Error bars represent ± SEM from two biological replicates. ***p<0.001, * p<0.05.

**Figure 5:**
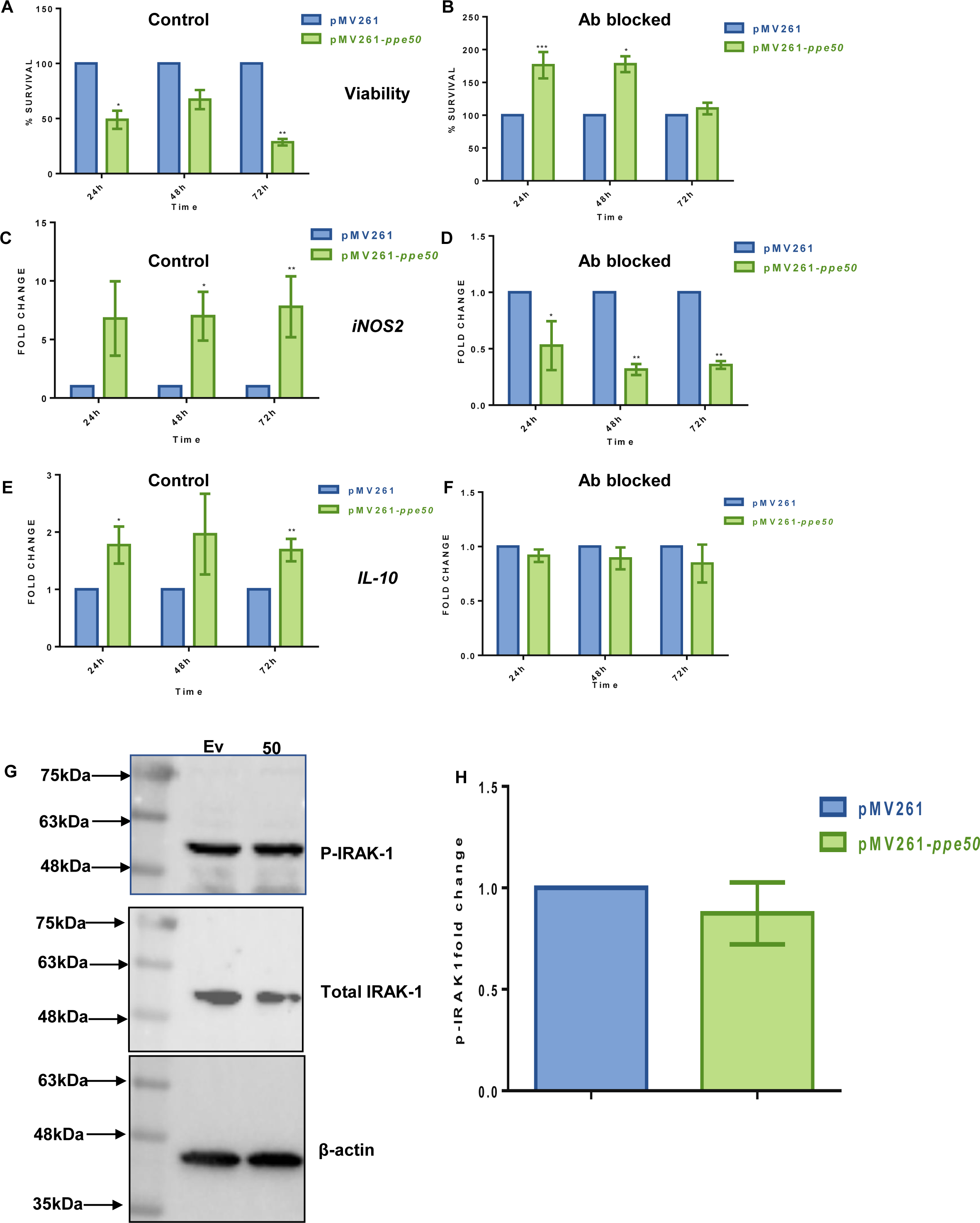
Effect of TLR1 blocking on CFU counts, iNOS*2* & *IL-10* transcript levels, and IRAK-1 phosphorylation. Growth index (A, B), *iNOS2* transcript levels (C, D), transcript levels of *IL-10* in THP-1 macrophages (E, F) infected with a recombinant *M. smegmatis* strain expressing the empty vector pMV261 (Ev) and *PPE50*, with and without antibody blocking. (G) Western blots showing levels of-actin levels in lysates from TLR1 blocked β infected macrophages (H) Densitometric quantitation of the blots in (G). The data represents two biological replicates. Error bars represent ± SEM from two biological replicates. ***p<0.001, **p<0.005, * p<0.05.

## DISCUSSION

While there is experimental evidence for the individual PE and PPE proteins, and PE proteins encoded in operons, in the modulation of host immunity and pathogenesis (20, 21, 33), there are no reports examining the involvement of the unusual PPE/PPE clusters in *M. tb* pathogenesis. Despite long-standing evidence for *PPE50* upregulation under conditions of hypoxia (7), and regulation of *PPE51* expression by the dormancy and hypoxia-induced DevRS TCS (2), the potential role of the *PPE50-PPE51* cluster in the pathophysiology of *M. tb* had not been investigated thus far.

Our RT-PCR analysis revealed that *PPE50* and *PPE51* are expressed as one transcriptional unit. Although this could have been predicted from the genomic coordinates with only 61 bp separating *PPE50* and *PPE51* (genomic coordinates 3501794-3502936), our data confirmed their operonic status. This observation was further validated by performing growth dependent expression analyses of these genes in *M.tb* H37Rv, where the two genes were seen to be coordinately expressed in all growth phases from log to stationary (Fig S8). Based on our experience of finding interaction between co-operonic members of this protein family (20, 21), we tested this possibility, and established using both genetic and biochemical approaches, that PPE50 and PPE51 indeed interact, the first record of a PPE-PPE heterodimeric pair. This opened up the possibility that the two proteins might play a role in *M. tb* physiology both independently as well as in a complex. To examine their possible role in mediating interactions at the mycobacterium-macrophage interface, we used the *M. smegmatis* model of macrophage infection that our laboratory has used extensively to interrogate the role of this family of proteins in host-pathogen interactions (20, 21, 34, 35). On infecting THP-1 macrophages with recombinant *M. smegmatis* expressing *PPE50*, *PPE51*, and the entire cluster, we observed that all these strains showed reduced survival compared to the control strain expressing the empty vector. We were surprised with this observation since the predominant phenotypes of recombinant *M. smegmatis* expressing the PE-PPE class of proteins in macrophages is that of enhanced survival compared to controls. Since this phenotype was reproducible and the reduced CFU counts correlated with an enhancement in *iNOS2* levels, we were convinced that this finding was genuine. We searched to see if any other such observation existed in the literature, and discovered that *M. smegmatis* expressing the stand alone genes *PPE10, PPE24*, showed low CFUs than their controls in RAW macrophages (31). However, in this report, the authors do not provide a corresponding measure of *iNOS2* levels for this data. Although unusual, we believe that this phenotype signifies that *M. tb* possesses the ability to reduce its infection load as a possible strategy for immune evasion.

In addition to reduction in bacillary viability, we also found that the anti-inflammatory cytokine IL-10 was consistently upregulated in macrophages infected with recombinant *M. smegmatis* expressing *PPE50* as a function of time, whereas strains expressing *PPE51* and the entire cluster showed none or inconsistent elevation in expression of *IL-10*. Since IL-10 inhibits the production of host-protective pro-inflammatory cytokines (31, 36, 37), the ability of PPE50 to induce an elevation in its levels makes it an important component of the anti-inflammatory arsenal of *M. tb*. Operating on the hypothesis that PPE50 mediates *IL-10* induction through ligand-receptor interaction, we performed receptor pull-down experiments and discovered TLR1 to be the cognate receptor for PPE50, with signal activation through the receptor being indicated by an increase in IRAK1 phosphorylation. In the signaling assay we also noted that IRAK1 phosphorylation was not induced in macrophages infected with recombinant *M. smegmatis* strains expressing either *PPE51*, or the entire operon. This observation, coupled with the lack of IL-10 upregulation in these very macrophages, led us to infer a negative regulatory role for PPE51 on PPE50 function when the two are in a complex (Fig. 4A). The reversal of all the phenotypes we observed on infection of THP-1 macrophages, on pre-treatment of macrophages with an anti-TLR1 antibody prior to infection, unequivocally demonstrated the functional outcome of PPE50-TLR1 interaction. This establishes PPE50 as a novel TLR1 ligand, and to the best of our knowledge only the second member of the PE-PPE family after PPE17 (38) that can bind and signal through this receptor.

Our findings suggest that the *PPE50-PPE51* cluster could play a role in facilitating bacillary persistence through decreasing of bacillary load and upregulation of the anti-inflammatory response, a two-pronged strategy to evade host immune surveillance. These observations strongly suggest that PPE50 and PPE51 are immunomodulatory proteins and are likely to participate in either the establishment or persistence of infection by hampering the mounting of immune response to *M. tb* infection *via* an altered TLR1 signaling pathway (Fig. 6). Performing macrophage and mouse infections with knock-out strains of *M. tb* carrying deletions in *PPE50*, *PPE51*, and the entire operon, would help solidify this hypothesis - these experiments are proposed in the next phase of this study.

**Figure 6:**
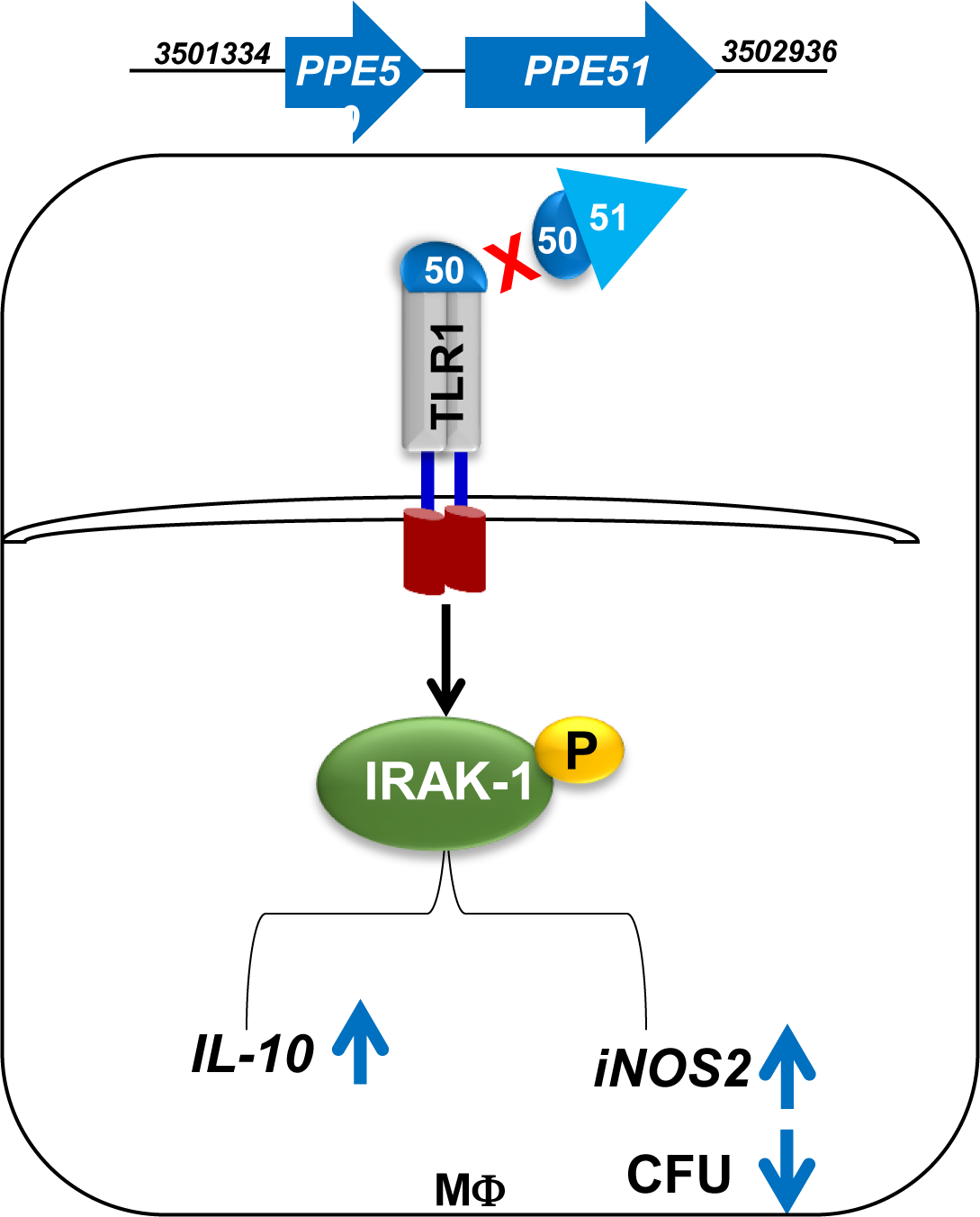
Model depicting the immunomodulatory/ immune evasion role of the *PPE50-PPE51* cluster in *M. tb* pathophysiology. MΦ - Macrophage.

Hypoxia, non-replicative persistence, thiol (diamide) stress, oxidative stress, iron deficiency, and starvation are all factors important to *M. tb* physiology, and have all been shown to affect the expression of different PE/PPE genes (2, 7). This suggests that the functions of these proteins are not redundant, but rather they serve distinct purposes during different stages of disease progression, depending on the pathogen’s niche. The pathogenic process is probably enhanced when these proteins and other PPE proteins operate together. It has been hypothesized that both cognate and non-cognate ESX-secreted PE/PPE complexes and their constituent proteins have virulence-effector roles (39). Recent reports have shown the YxxxD/E motif to be a conserved secretion signal, which has been found in all known substrates or substrate complexes of the mycobacterial type VII secretion system (40). Since this signature is conserved in PPE51 as well, the localization/ export of these proteins could be tracked in *M. tb* strains with mutations in the *esx* genes to test this theory.

In summary, this study represents the first demonstration of an independent immunomodulatory function of a PPE-PPE protein cluster and strongly suggests that the *PPE50-PPE51* cluster could potentially play a pivotal role in the establishment and persistence of *M. tb* infection in the host.

## Author contributions

T. R. R. designed the study. R. P. M., K. P. and S. Y. performed the experiments. T. R. R. and R.P.M. analysed the data. T.R.R. and R. P. M wrote the manuscript.

