## Supplementary Material - 8 Supplementary Figures and 1 Supplementary Table for "Delineating the functional role of the *PPE50 (Rv3135) - PPE51 (Rv3136)* gene cluster in the pathophysiology of *Mycobacterium tuberculosis*"

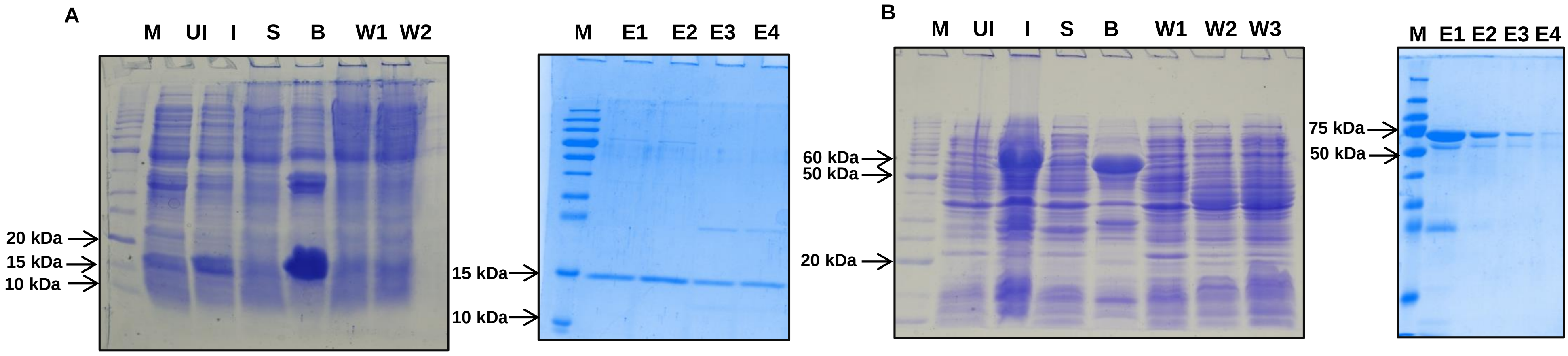


**Figure S1 : Expression and purification profile of PPE50, PPE51** (A) SDS-PAGE profiles showing expression and purification of CT-6xHIS-tagged PPE50. (B) ) SDS-PAGE profiles showing expression and purification of NT-GST tagged PPE51. UI - Uninduced sample, I - Induced sample, S - Supernatant, B - Bound Beads, M - Marker, W - Wash, E - Eluate.

A

| | Input control | | | Pull down | | | |
|---|---|---|---|---|---|---|---|
| PPE50-6xHis | - | + | - | - | + | + | + |
| GST-PPE 51 | + | - | - | + | - | - | + |
| GST | - | - | + | - | - | + | - |
| Protein A Agarose | - | - | - | + | + | + | + |

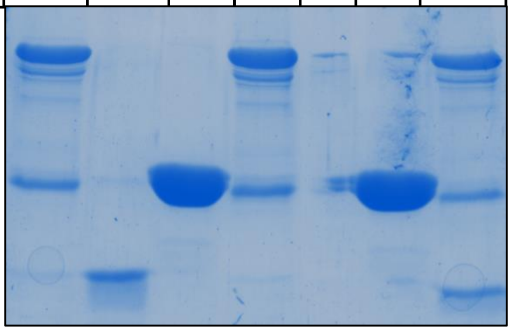

B

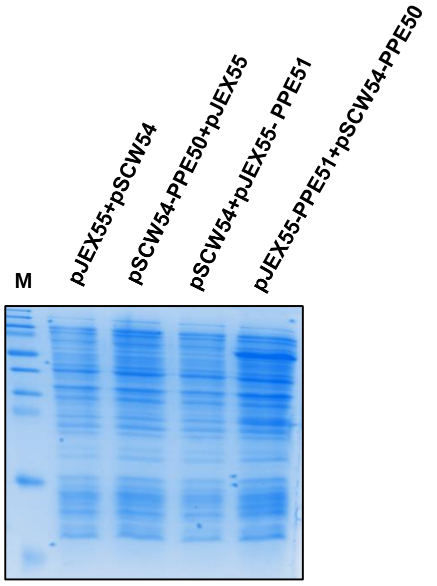


**Figure S2: SDS-PAGE profiles demonstrating equivalent loading for PPE50-PPE51 interaction assays** (A) Samples used for *in vitro* co-IP reactions of PPE50-6xHis and GST-PPE51 were pulled down with an anti-GST antibody and probed with an anti-His antibody. (B) Samples used for co-IP reactions of PPE50-PPE51 from *M. smegmatis* expressing PPE51-cmyc and PPE50-6xHis.

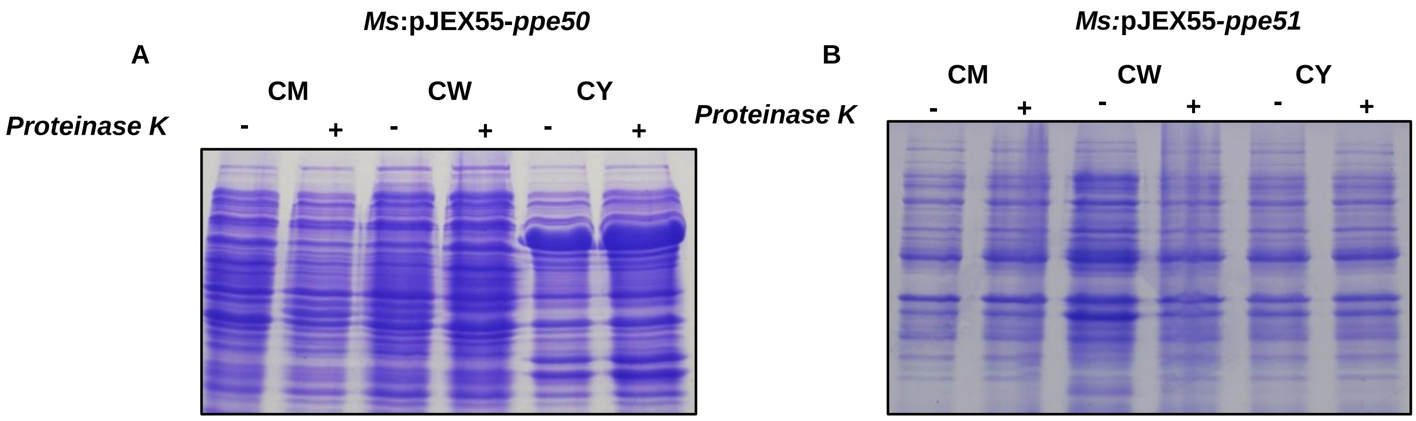


**Figure S3 : Sub-cellular localization and surface accessibility**
CM - cell membrane fraction, CW - cell wall fraction, CY - cytoplasmic fraction. Coomassie stained SDS PAGE profiles showing equal loading for the sub-cellular fractions of Proteinase K treated *M. smegmatis* expressing PPE50 (A) and PPE51 (B).

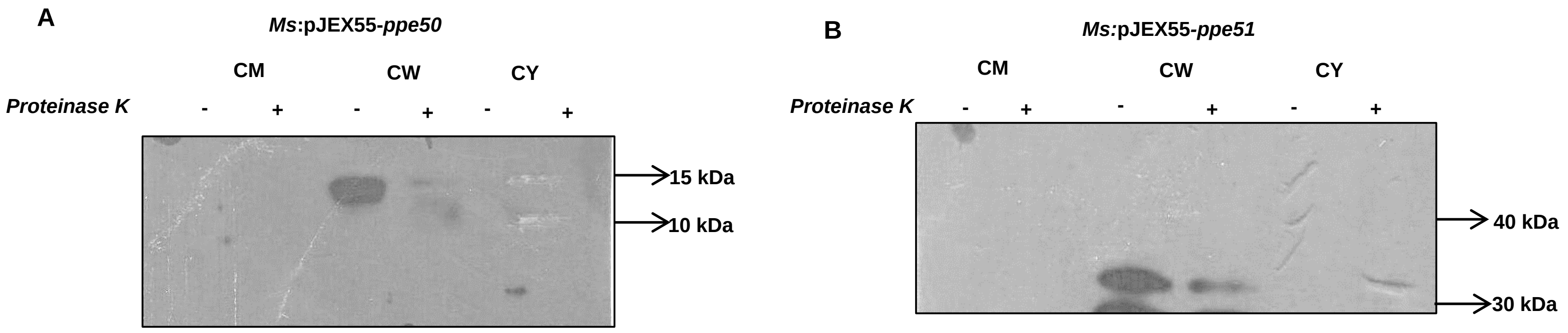


**Figure S4 : Sub-cellular localization and surface accessibility**. Immuno-detection of PPE50 (A) and PPE51 (B) in subcellular fractions of Proteinase K treated recombinant *M. smegmatis* strains expressing PPE50-myc and PPE51-myc. All proteins were detected using an anti c-myc mAb. CM - cell membrane fraction, CW - cell wall fraction, CY - cytoplasmic fraction.

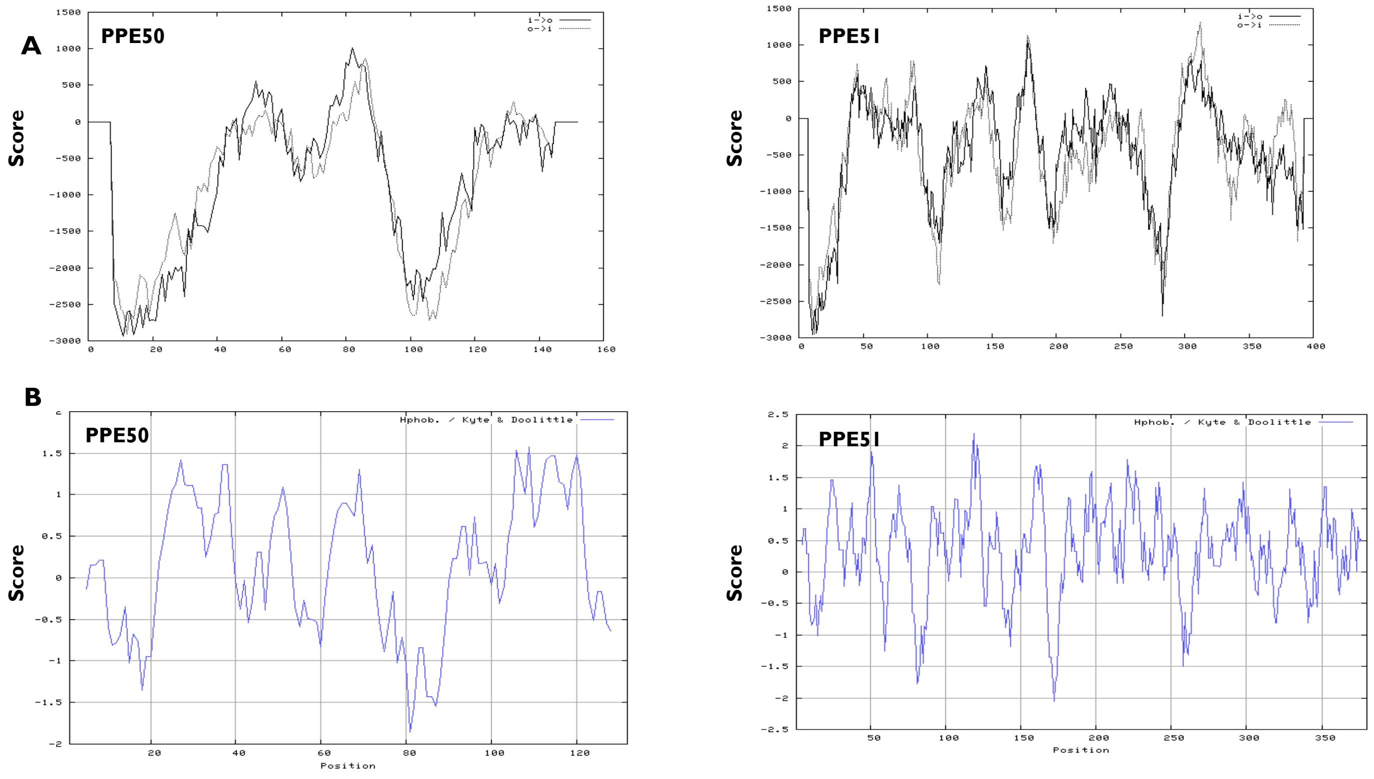


**Figure S5: *in silico* protein sequence analysis of *M. tb* PPE50 and PPE51**. Transmembrane prediction (A), and Hydrophobicity (B) analyses of PPE50 and PPE51.

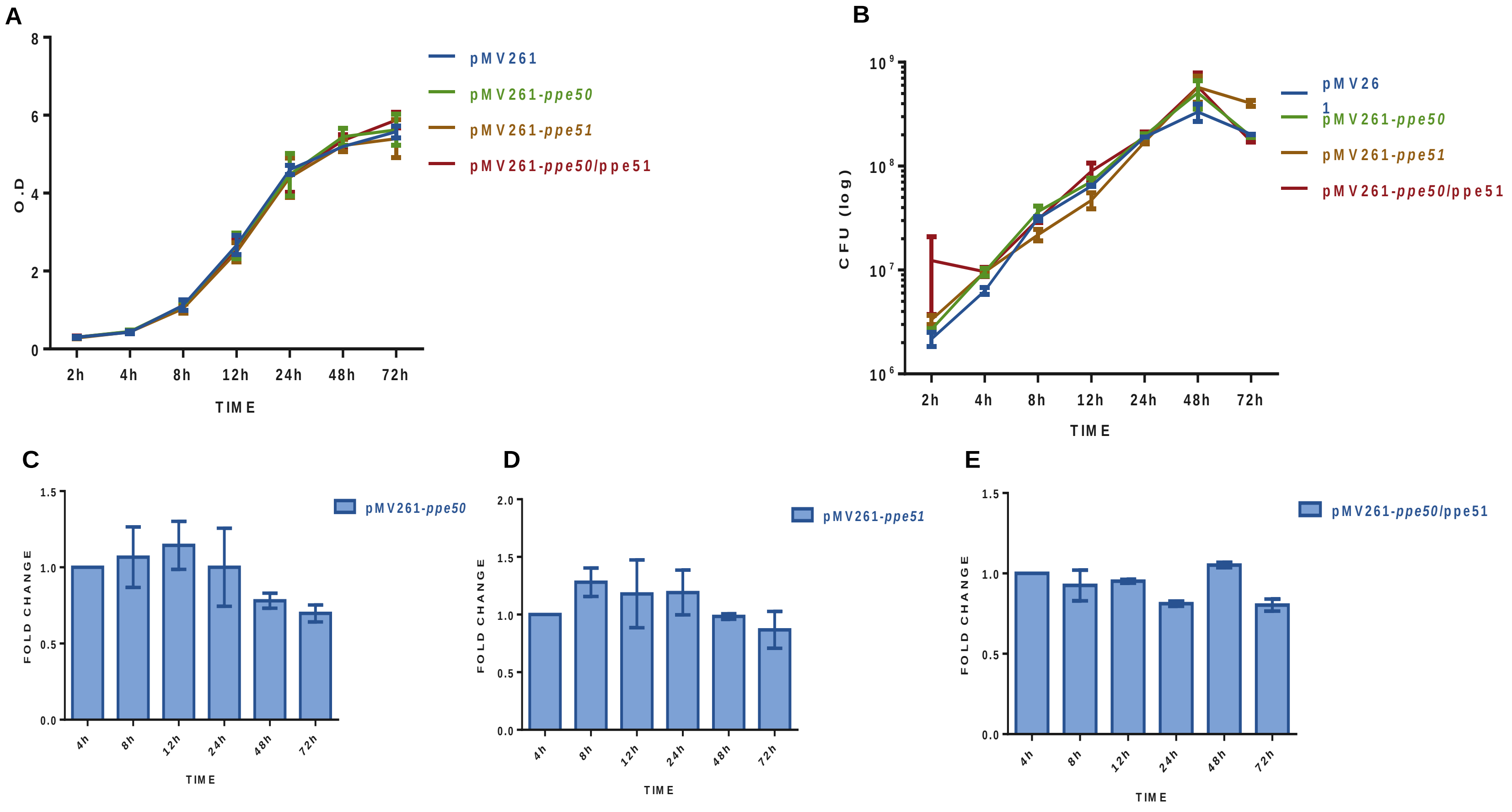


**Figure S6: Growth and gene expression analysis of recombinant *M. smegmatis* expressing *PPE50* and *PPE51*** *In vitro* growth profiles of *M. smegmatis* expressing pMV261, *PPE50*, *PPE51*, and *PPE50-PPE51* were obtained by measuring OD (A) and estimating CFU counts (B). Real-time RT-PCR quantification of *PPE50* (C) and *PPE51* (D) transcripts, as well as *PPE50-PPE51* (E) transcripts, as a function of growth. Transcript levels are shown relative to the homologous gene's mRNA levels at 4h, which is given a value of 1. The error bars represent ± SEM.

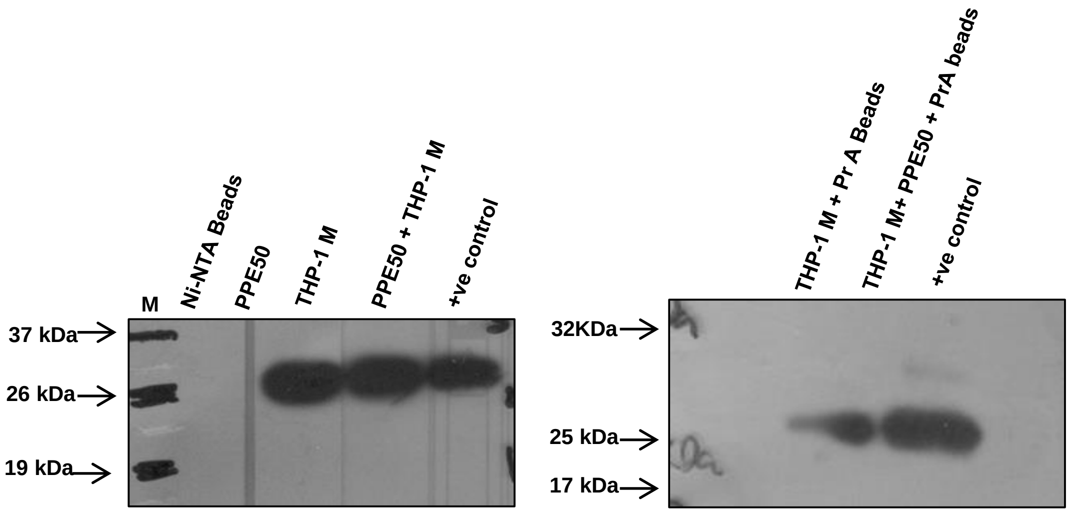


**Figure S7: Western blots showing equal loading for the pull-down (A) and reverse pull down (B) experiments shown in Fig. 4A, B.** The listed samples were separated by SDS-PAGE and Western blotting was performed with an antibody to CD81

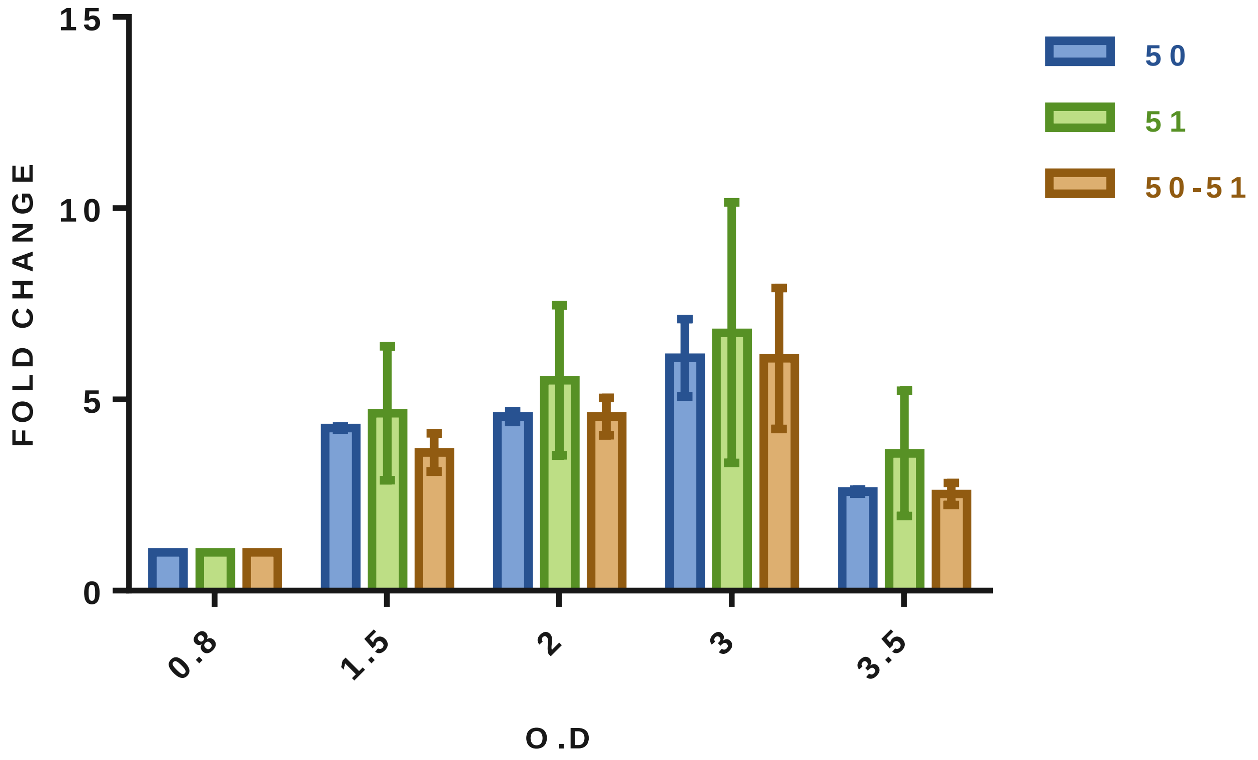


**Figure S8: Gene expression profiles of *PPE50* and *PPE51 in M. tb* H37Rv** Real-time RT-PCR quantification of *PPE50, PPE51* and *PPE50-PPE51* transcripts. Transcript levels are shown relative to the cognate gene's mRNA levels at 0.8 OD, which is given a value of 1. The error bars represent ± SEM

**Table S1: Primers used in this study**

| **Name** | **Description** | **Sequence** |
|---|---|---|
| pMV261- PPE50-F | FP (Forward Primer) for cloning *PPE50* in pMV261 | 5’CGCGGATCCATGGACTA CGCGTTCTTACCAC3’ |
| pMV261- PPE50-R | RP (Reverse Primer) for cloning *PE50* in pMV261 | 5’CCCAAGCTTTCAAGGTGG AGTGCCAGCG3’ |
| pMV261-PPE51-F | FP for cloning *PPE51* in pMV261 | 5’CGCGGATCCATGGATTTCGC TGTTACCACC3’ |
| pMV261-PPE51-R | RP for cloning *PPE51* in pMV261 | 5’CCCAAGCTTCCACCC GCGGCAGGGTAA3’ |
| pJEX55-PPE50-F | FP for cloning *PPE50* in pJEX55 | 5’CGCGGATCCATGGACTA CGCGTTCTTACCAC3’ |
| pJEX55-PPE50-R | RP for cloning *PPE50* in pJEX55 | 5’CCCAAGCTTTCAAGGTGG AGTGCCAGCG3’ |
| pJEX55-PPE51-F | FP for cloning *PPE51* in pJEX55 | 5’CGCGGATCCATGGATTTC GCACTGTTACCACC3’ |
| pJEX55-PPE51-R | RP for cloning *PPE51* in pJEX55 | 5’CCCAAGCTTCCACCCGC GGCAGGGTAA3’ |
| pET22b-PPE50-F | FP for cloning *PPE50* in pET22b | 5’CCATGCCTCGAGGCCGC CCACGACCCCGTAC3’ |
| pET22b-PPE50-R | RP for cloning *PPE50* in pET22b | 5’CCGCTCGAGCGGCTCGA GAGGTGGAGTGCCAGCGG TGT3’ |
| pGEX6p1- PPE 51F | FP for cloning *PPE51* in pGEX6p1 | 5’CGCGGATCCATGGATTTC GCACTGTTACCACC3’ |
| pGEX6p1- PPE51R | RP for cloning *PPE51* in pGEX6p1 | 5’ATAAGAATGCGGCCGCTT AGTGGTGGTGGTGGTGGT GCCCTGCCGCGGGTGGGT 3’ |
| RT-PPE50 F | FP for expression analysis of *PPE50* in *M. smegmatis* (RT PCR) and co-operonic analysis | 5’GCGCGTATGTACAGCGG T3’ |

| | | |
|---|---|---|
| RT-PPE50 R | RP for expression analysis of *PPE50* in *M. smegmatis* and co-operonic analysis | 5’GTCCGCTCCAGCCATTCC 3’ |
| RT-PPE51 F | FP for expression analysis of *PPE51* in *M. smegmatis* and co-operonic analysis | 5’GCTGACGATTCCGAGCTT CA3’ |
| RT-PPE51 R | RP for expression analysis of *PPE51* in *M. smegmatis* and co-operonic analysis | 5’AACTCGCCGATCCCAAAG TC3’ |
| PPE50- JF | FP for co-expression analysis of *PPE50-PPE51* junction in *M.tb* and co-operonic analysis | 5’CGGCATTCGAGCAGGCT C3’ |
| PPE50-JR | RP for co-expression analysis of *PPE50-PPE 51*junction in *M.tb* and co-operonic analysis | 5’GTAACAGTGCGAAATCCA TATTGT3’ |
| pUAB400-PPE50F | FP for MPFC analysis of PPE50-PPE51 | 5'ACAAGAATTCATGGACTA CGCGTTCTTACCACCGG3’ |
| pUAB400-PPE50R | RP for MPFC analysis of PPE50-PPE51 | 5’TAGGAAGCTTTCAAGGTG GAGTGCCAGCG3’ |
| pUAB300-PPE51F | FP for MPFC analysis of PPE50-PPE51 | 5’CATCGGATCCATGGATTT CGCACTGTTACCACC3’ |
| pUAB300+PPE51R | RP for MPFC analysis of PPE50-PPE51 | 5'GAACAAGCTTTTACCCTG CCGCGGGTGGGT3' |
